# TRACE: An unbiased method to permanently tag transiently activated inputs

**DOI:** 10.1101/2020.02.02.927194

**Authors:** Nathalie Krauth, Valentina Khalil, Meet Jariwala, Noëmie Mermet-Joret, Anne-Katrine Vestergaard, Marco Capogna, Sadegh Nabavi

## Abstract

A fundamental interest in circuit analysis is to parse out the synaptic inputs underlying a behavioral experience. Toward this aim, we have devised an unbiased strategy that specifically labels the afferent inputs that are activated by a defined stimulus in an activity-dependent manner. We validated this strategy in four brain areas receiving known sensory inputs. This strategy, as demonstrated here, accurately identifies these inputs.

## Introduction

Most of the brain regions receive a large number of neuronal inputs; this poses a challenge to circuit analysis since only a fraction of these inputs conveys information for a defined behavior. Identifying these inputs is crucial to gaining an insight into the neural circuits that underlie the behavior. Currently, there is no direct way of achieving this goal. A common strategy is to use retrograde tracing viruses which are designed to identify all the inputs to the region of interest^1–3^. To identify the specific inputs, the researcher must then rely on a combination of trial and error, an educated guess, and previous findings. A more efficient way would be to label only the inputs that are activated by the stimuli. Recent developments in the use of immediate early gene promoters offer such an opportunity^4,5^. The underlying mechanism is simple: a gene of interest is expressed under the control of an activity-dependent promoter such as Arc or c-fos. The neurons that are activated by the behavioral experience will express the gene of interest such as a fluorescent marker. This approach, as currently used, however, does not reveal the active inputs.

Here, we introduce a novel approach, Tracing Retrogradely the Activated Cell Ensemble (TRACE), which selectively labels the afferent inputs that are activated by a defined stimulus and project to the region of interest. It combines activity-dependent labeling with virus-mediated retrograde tracing. This approach is unbiased, as it does not rely on pre-existing knowledge of candidate regions and offers high temporal (minutes) and spatial (cellular scale) resolution.

## Results

TRACE is based on two recently developed methods: 1) labeling of active neurons wherein neurons express the tamoxifen-inducible CreER-recombinase under the control of an activity-dependent promoter such as Arc or c-fos^4,6,7^, and 2) labeling with a retrograde virus, such as retroAAV, that expresses a marker gene in a recombination-dependent manner^3^. In this work, we chose retroAAV as it is one of the most efficient retrograde viruses available showing little toxicity. Importantly, TRACE can be adapted to any DNA-based retrograde virus. TRACE works as following: first, a retroAAV carrying a Cre-dependent marker is injected into a target brain area. Then, the virus is taken up by post-synaptic neurons in the target region as well as by the axons of the projecting neurons, but the neurons expressing the virus remain unlabeled in the absence of tamoxifen and neuronal activity. As the animals are exposed to a behavioral experience, only the projecting neurons activated in the short time period of the behavior express Cre-ER. We then inject 4-hydroxytamoxifen (4-OHT) to induce CreER recombinase translocation into the nucleus and recombination of the marker genes. This results in a permanent expression of the marker genes in active neurons projecting to the target area (Fig. 1a). To demonstrate the input specificity of this method, we used this approach in four different behaviorally-relevant neuronal circuits.

**Fig. 1.**
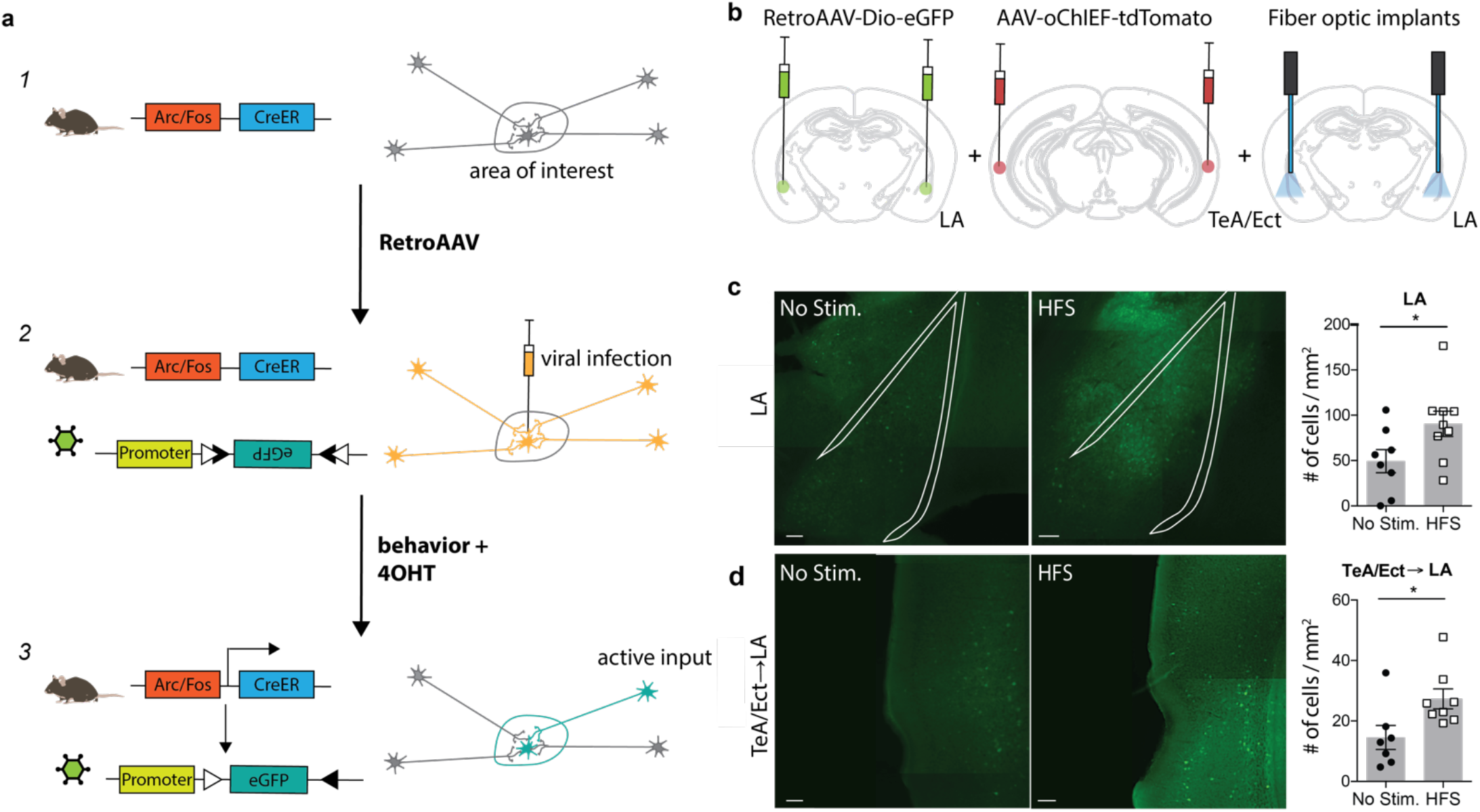
TRACE method. TRACE labeling of afferent cortical inputs to lateral amygdala after high frequency stimulation (HFS). **a**, Schematic of the TRACE method: (1) A transgenic mouse expresses the tamoxifen-inducible CreER-recombinase under the arc or c-fos promoter. (2) The retroAAV infects the cells in the area of interest and is also taken up by cellular projections of other areas to this region. The retroAAV has a floxed eGFP, which will not be expressed in this state. (3) Upon 4-OHT injection the CreER recombination occurs in active cells. Consequently, eGFP is expressed in all active cells that were previously infected with the virus. This causes labeling of active cells in the area of interest as well as in their respective inputs. **b**, Experimental schematic. RetroAAV-Dio-eGFP was injected into the lateral amygdala (LA), while the TeA (temporal association area)/Ect (ectorhinal cortex) were injected with the channelrhodopsin carrying virus AAV-oChIEF-tdTomato. The optic fibers were placed above the LA in order to activate the channelrhodopsin infected projections from TeA/Ect to the LA. 4-OHT was injected 2 hours after HFS. The quantification of active labeling in the LA and TeA/Ect was performed 1 week after the HFS treatment. **c**, Exemplary images of the LA, outlined with the white line, in mice receiving HFS or no stimulation. Green shows activity-mediated eGFP fluorescent expression. Graph (right panel) shows quantification of the fluorescently-tagged neurons. Labelling in the LA is significantly higher in the optically-activated animals (unstimulated group n = 8, mean ± SEM, HFS group n = 9; p-value = 0.0474(unpaired t-test); scale bar: 100um). **d**, Representative images of the TeA/Ect, outlined with the white line, in mice receiving HFS or no stimulation. Green shows activity-mediated eGFP fluorescent expression in the LA-projecting neurons within TeA/Ect. Graph shows quantification of the fluorescently-tagged neurons. Activity-dependent retrograde labeling is significantly higher in the activated projections from TeA/Ect to LA (unstimulated group n = 7, HFS group n = 8; p-value 0.014 (Mann-Whitney test); scale bar: 100um).

### TRACE labeling of afferent cortical inputs to lateral amygdala after high frequency stimulation (HFS)

First, we chose to apply our method to label afferent inputs from the temporal association area and ectorhinal cortex (TeA/Ect) to the lateral amygdala (LA), since this is a well-characterized pathway with clear-cut behavioral significance^8^. We injected an AAV expressing a variant of the light-activated channel ChR2, oChIEF, into the TeA/Ect and a retroAAV virus expressing eGFP in a Cre-dependent manner in the LA of ArcCreER mice. A fiber optic was placed above the LA to deliver optical stimulation (Fig. 1b and Supplementary Fig. 1a). Three to four weeks after the viral injection, multisensory inputs to the LA were stimulated with high-frequency light pulses, followed by injection of 4-OHT. We observed significantly more eGFP-tagged neurons in the post-synaptic neurons in the LA (Fig. 1c) and, more importantly, in presynaptic neurons in the TeA/Ect of the mice receiving optical activation compared to non-stimulated mice (Fig. 1d).

### TRACE identifies inputs activated by aversive experience

Next, we explored whether TRACE could identify the inputs for an aversive foot-shock. We chose this particular stimulus, since it has been widely used in studies on associative learning and valence experiences^9^. For this, we focused on the pathway from the medial prefrontal cortex (mPFC) to the lateral habenula (LHb) and to the nucleus accumbens (Nacc), since these inputs are known to convey an aversive signal for a foot-shock to these targeted areas^9,10^. We injected retroAAV viruses expressing two different fluorescent markers in a Cre-dependent manner in the LHb (retroAAV-Dio-eGFP) and in the Nacc (retroAAV-Dio-mCherry) of cfosCreER mice (Fig. 2a and Supplementary Fig. 1b). Two weeks after the virus injection, mice were divided into two groups. The test group received a series of foot-shocks followed by injection of 4-OHT, whereas the control group received 4-OHT in their home cages. Animals receiving foot-shocks had significantly higher fluorescent marker expression in the mPFC compared to the control group (Fig. 2b,c), consistent with the previous reports on the aversive nature of the mPFC inputs to the LHb and Nacc^9,10^.

**Fig. 2.**
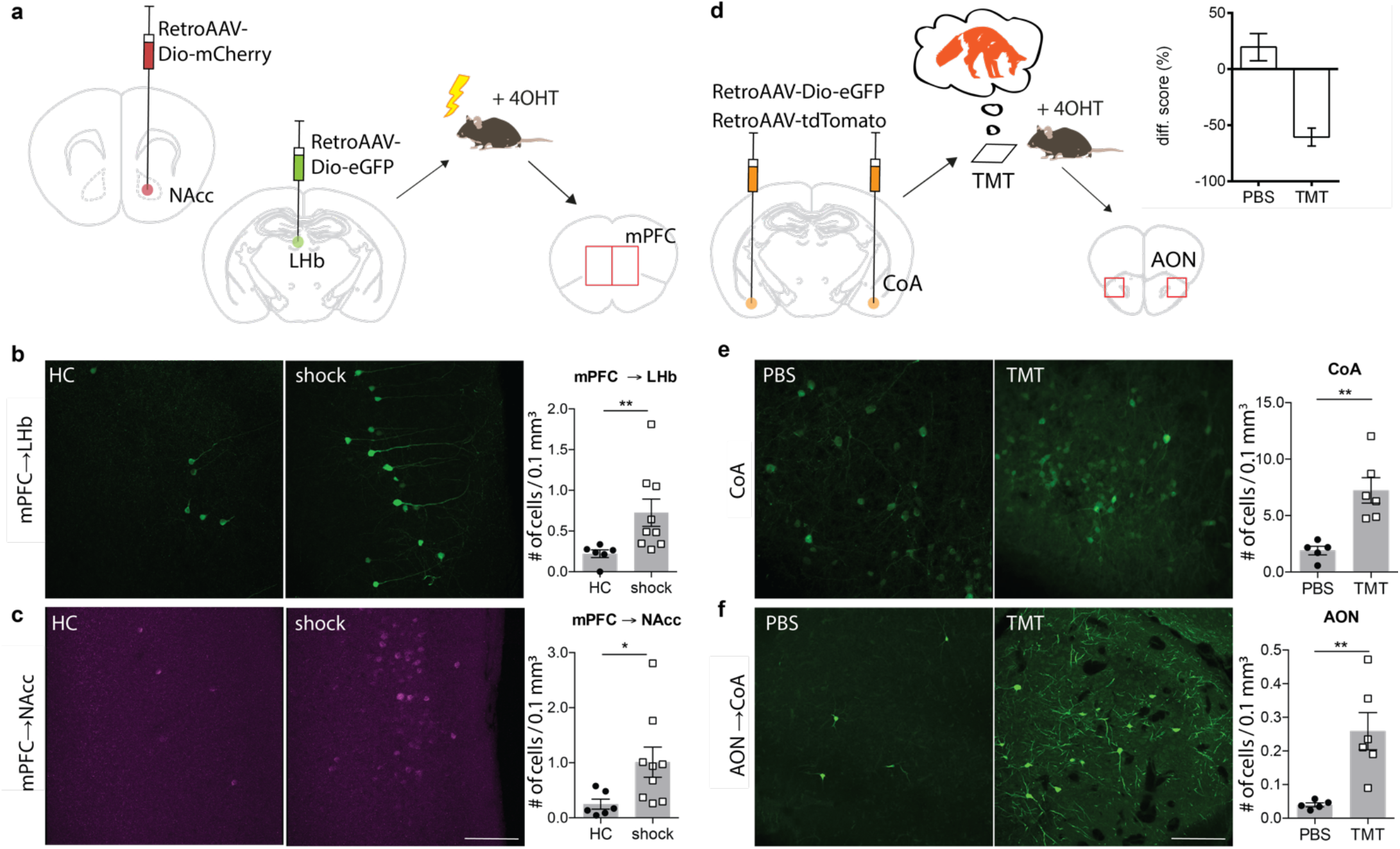
TRACE identifies inputs activated by aversive experiences. **a**, Experimental schematic. Injection of RetroAAV-Dio-eGFP into the left hemisphere LHb. Injection of RetroAAV-Dio-mCherry into the right hemisphere NAcc. Two weeks after virus infection of LHb and NAcc, a group of animals was exposed to 20 mild electric foot-shocks within a 10-minute session while a control group remained in their home cages. 4-OHT was injected two hours after testing. The evaluation of active labeling in LHb, NAcc and mPFC in both hemispheres was performed 1 week after testing. **b**, Magnified images of the mPFC in mice receiving foot-shock. Green shows activity-mediated eGFP fluorescent expression in the LHb-projecting neurons within mPFC. As graph shows mean ± SEM), TRACE-mediated labeling in the LHb-projecting neurons in the mPFC of animals exposed to the foot-shocks is significantly higher than the home cage group (home cage group n = 6, shock group n = 9; p-value= 0.0016 (Mann-Whitney test); scale bar: 100 um). **c**, Same as b, for the NAcc-projecting neurons in the mPFC (home cage group n = 6, shock group n = 9; p-value = 0.012 (Mann-Whitney test). Color-converted magenta depicts the activity-mediated mCherry fluorescent expression. **d**, Experimental schematic. Injection of RetroAAV-Dio-eGFP and RetroAAV-tdTomato into the cortical amygdala. Three weeks after virus infection of CoA, the animals were exposed to the 2,3,5-Trimethyl-3-thiazoline (TMT) odor, followed by 4-OHT injection two hours later. Evaluation of active labeling in CoA and AON was performed 2 weeks later. The graph shows the time spent within the chamber (% of total time) comparing habituation vs. test sessions for both PBS and TMT groups (PBS group n=7; TMT group n = 9; 0.0003 (Mann-Whitney test). e, Magnified images of the CoA in mice exposed to PBS or TMT. As shown in the right graph, activity-dependent eGFP expression is significantly higher in TMT-exposed mice (PBS exposed n = 5, TMT exposed n = 6; p-value = 0.0043 (Mann-Whitney test); scale bars: 100 um). f, Magnified images of the anterior olfactory nucleus AON) in mice exposed to PBS or TMT. TRACE-mediated labeling in the CoA-projecting neurons in the AON of animals exposed to TMT is significantly higher (PBS group n = 5, TMT group n = 6; p-value= 0.0043 (Mann-Whitney test)).

### TRACE identifies inputs activated by innate aversive experience

Finally, we chose to characterize inputs onto the cortical amygdala (CoA) driven by an innate odor-stimulus, which is likely to recruit long-range projections from the olfactory nuclei^11^. As a stimulus, we used 2,3,5-trimethyl-3-thiazoline (TMT), a volatile component found in fox secretions, that is known to activate the inputs from the olfactory cortex to the CoA^12^. The odor-driven activation of these inputs to the CoA induces an innate avoidance behavior (Fig. 2d and Supplementary Fig. 2). We injected retroAAV expressing Cre-dependent eGFP in the CoA (Fig. 2d and Supplementary Fig. 1c). Three weeks after the virus injection, mice were divided into two groups. The test group were exposed to TMT, and subsequently they received a dose of 4-OHT. The control group received the same treatment as the test, except that TMT was replaced with phosphate buffer saline (PBS). In the test group, we observed significantly more eGFP expressing cells in the anterior olfactory nucleus (AON) than in the control group (Fig. 2e,f). This is consistent with previous reports showing a TMT-induced activation of the AON^13^. To examine the input specificity of TRACE, we quantified eGFP-labelled cells in selected regions, which send pronounced projections to the CoA, as characterized by our activity-independent retroAAV labelling. However, we did not observe a difference between the control and the test groups (Supplementary Fig. 3). This indicated that labelling is specific to the activated regions and merely projecting to a target region is not sufficient for eGFP labelling.

### Discussion and Outlook

Considering the complexity of the circuits underlying behaviors, unbiased approaches are particularly valuable in identifying the inputs which drive behavioral responses. The applications described above show that TRACE can identify active afferent inputs with virtually no need for a priori knowledge of their origins. Our approach, in principle, can be combined with genetically-encoded indicators such as GCaMP to monitor the activity of input regions for different behavioral tasks. Also, TRACE can be used to deliver chemo- and opto-genetic tools to the functionally relevant input neurons for further circuit manipulations. Since TRACE is a non-transsynaptic tracer, it is well suited for identifying neuromodulatory inputs which convey their signals through volume transmission rather than direct synaptic connections^14^. A potential concern in using a non-transsynaptic tracer as TRACE could be represented by the infection of axons passing through the injection site. However, there has not been such report for retroAAV.

In using TRACE, the same considerations must be taken as those for activity-dependent promoters and virus-mediated labelling. Viral tropism has been a recurring concern in investigating circuit mapping^3,15^. New strategies such as receptor complementation have been introduced to overcome the problem with tropism^16^. Alternatively, retroAAV virus can be complemented with a retrograde virus of different tropism, such as canine adenovirus type-2. In a similar way, activity-dependent promoters display region specificity^4,7^, as TRACE was not efficient in labelling the olfactory bulb as a source for aversive input to the CoA despite its role in odor-driven innate aversive behavior^11^. However, with the rapid development in activity-dependent labeling systems, we are closer than ever to gain brain-wide access to neurons activated by a particular experience^4,6,7,17,18^. Despite these considerations, we anticipate that TRACE, as it stands, will be generalizable and complementary to other circuit analysis methods for elucidating how neuronal activity in connected ensembles drives complex behaviors.

## Author Contributions

The project was designed by SN and NK. The manuscript was written by SN, MC, and NK. Figures were designed and composed by NK, NMJ and VK. For HFS and TMT testing NK and NMJ performed the surgeries. For the foot-shock experiments surgeries were done by NK. VK performed the TMT behavior. NK and MJ performed the HFS behavior. NK, MJ and AKV performed the foots-hock behavior. Imaging for HFS and foot-shock was done by NK and MJ. Imaging for TMT experiments was done by VK and NMJ. Cell counting was done by VK, MJ and NK. Statistical analysis and graphs were done by NK and VK. SN and MC supervised the research. All authors discussed the results and contributed to the revision of the manuscript.

## Acknowledgements

We thank the members of the Nabavi lab, in particular Islam Faress for suggestions. This study was supported by ERC grant 679714 and Lundbeck foundation grant to SN and the Danish National Research Foundation grant DNRF133 to SN and MC.

## Competing Interests

The authors declare no competing interests.

Correspondence and requests for materials should be addressed to SN

## Methods

### Animals

ArcCreER heterozygous mice B6.Cg-Tg (Arc-cre/ERT2)MRhn/CdnyJ (JAX stock #022357) from Jackson labs were used in the optical activation experiments and the odor driven innate fear experiments. In the foot-shock experiments c-fosCreER heterozygous mice B6.129 (Cg)-Fostm1.1 (cre/ERT2)Luo/J (JAX stock #021882) from Jackson labs were used.

All mice were given food and water ad libitum and were kept at a 12h light/dark cycle, with the light being on during daytime. All behavioral tests were performed during day-time, however the animals were kept in isolation and in a dark room 24 hours before and 72 hours after the testing days. Odor driven innate fear experiments were only performed with male mice, since it was observed that the female mice did not react to TMT in the same manner as their male littermates (data not shown). Both the optogenetic activation and shock-induced circuit experiments were performed with mice of both sexes and mixed test groups, as no difference was observed. All procedures involving animals were approved by the Danish Animal Experiment Inspectorate.

### Viral vectors and tracers

The AAV were obtained from the viral vector core facility at the University of Zurich in Switzerland. Virus titers were the following: 4.5×10^12^ for AAV-hSyn1-dlox-eGFP (rev)-dlox-WPRE-hGHp (A) (serotype AAV2 retro); 5.0×10^12^ for AAV-hSyn-oChIEF-tdTomato (Serotype AAV8); 1.0×10^13^ for AAV-hSyn-oChIEF-td tomato (serotype AAV2 retro); 4.8×10^12^ for AAV-hSyn1-dlox-mCherry(rev)-dlox-WPRE-hGHp (A) (serotype AAV2 retro); 7.0×10^12^ for AAV-hSyn1-dlox-mCherry (rev)-dlox-WPRE-hGHp (A) (serotype AAV9).

### Stereotactic surgeries

Mouse surgeries were performed on 5 week-old mice to ensure low background labeling. The mice were anesthetized using a mix of 0.05 mg/ml of Fentanyl ((Hamel, 007007) 0.05 mg/kg), plus 5 mg/ml of Midazolam ((Hameln, 002124) 5 mg/kg) and 1 mg/ml of Medetomidine ((VM Pharma, 087896) 0.5 mg/kg) (FMM) and surgeries were performed using a stereotaxic frame (Kopf Instruments). The scalp was opened using scissors and holes were drilled using a Foredom high speed drill (Foredom Electric Co, K.1070-22). Coordinates were normalized to a bregma-lambda distance of 4.21 mm. In the case of viral injections, the animals were injected with a pulled 1 mm glass pipette using a picospritzer or Nanoject III Drummond) containing the respective virus mix for each experiment. 0.5 ul of virus mixtures were injected per location unless otherwise stated in the main text.

Viral injection coordinates for the optogenetic stimulation experiments were: LA from Bregma: anterior-posterior (AP), −1.6 mm; medio-lateral (ML), 3.45 mm; and dorsal-ventral (DV) from the skull, −4.0 mm and TeA/Ect AP, −2.5 mm and −3.0 mm; ML, 4.3 mm; DV, −2.0 and −2.4mm). The fiber implant (0.22 numerical aperture (NA), 200 um core diameter) was placed above the LA (AP, −1.5 mm; ML, 3.35 mm; and DV, −3.9 mm). Viral injection coordinates for the odor-activated innate fear experiments were: CoA (AP, −1.6 mm and −1.7 mm; ML, 2.7 mm; DV, −5.8 mm). Viral injection coordinates for the shock activated experiments were: LHb (AP, −1.45 mm; ML, 0.3 mm; DV, −2.8 mm) and NAcc (AP, +1.5 mm; ML, 1.0 mm; DV, −4.5 mm).

After the surgery, the wound was sutured and the mice received 200 ul of a local anesthetic (lidocaine 2%) to minimize pain from the surgery. The mice were given an antidote mix of 0.4 mg/ml Naloxone ((B. Braun, 115241) 1.2 mg/kg), plus 5 mg/ml of Atipamelozone Hydrochloride (2.5 mg/kg) and 0.5 mg/ml of Flumazenil Hameln, 036259) 0.5 mg/kg) to reverse the anesthesia and the animals were allowed to recover on a heating pad. Buprenorphine (Temgesic (Indivior UK Limited, 521634) 0.3 mg/ml) was added to the drinking water as an analgesic in the two days following surgery. The mice were allowed to recover for 2 weeks before behavioral tasks. The animals were checked daily after surgery.

All the viral injection sites and the optic fiber implants were confirmed histologically and the animals were excluded when the injection sites or the optic fiber implantation were misplaced.

### Behavioral testing

#### Optogenetic stimulation of LA

ArcCreER mice were habituated to the plugging of the laser patch cord for 5 days while exploring the testing cage for 10 mins. Following this training, the mice were intraperitoneally i.p.) injected with saline daily to habituate them for tamoxifen injections. One day before the testing day, the mice were moved to the dark housing, where they were individually housed until 72 hours after the testing. Optogenetic stimulation was performed in the home cages using a laser with a 450 nm wavelength (Doric lenses) and a dual fiber optic patch cord (Doric Lenses, 0.22NA, 200um core diameter). To induce antidromic spikes, we used a high-powered laser stimulation protocol^19^ consisting of 10 trains of light (each train having 100 pulses of light, 5 ms each, at 50-80 mW) at 90-s inter-train intervals controlled with a pulse generator (Pulse Pal, Open Ephys). IP injection of 4-OHT (10 mg/kg) was performed 2 hours after the behavior. 4-OHT was prepared following Ye et al. 2012.

#### Odor driven innate fear

ArcCreER mice were habituated to the context consisting in an open field arena (45 cm x 35cm) containing a small dark chamber (16.5 cm x 10 cm) in the center. The dark chamber had a single opening on the side (Genné-Bacon et al., 2016). Seven sessions of habituation were performed in 3.5 days, twice a day, and each session lasted 10 minutes (performed during day light). At the beginning of each session, the mice were placed randomly around the dark chamber. Between the morning session and the afternoon session, the mice were habituated to intraperitoneal injections. One day before the testing, the mice were moved to the dark housing, where they were kept individually housed until 72 hours after the testing. For both PBS and 2,4,5-Trimethyl-thiazoline (TMT) (Sigma), 20ul of the solution was dropped onto a filter paper fixed in the middle of the dark chamber. The TMT had previously been diluted 1:1 with PBS to lower the strength. The test group was exposed to PBS in the dark chamber in the morning and to TMT in the afternoon. The control group was exposed to PBS during both test sessions. The mice were positioned in front of the entrance of the dark chamber for the testing sessions. IP injection of 4-OHT (10 mg/kg) was performed 2 hours after TMT exposure. The behavior was recorded and analyzed manually with a video tracking system (ANY-maze).

#### Shock-activated circuit

cfosCreER mice were habituated to the context with 5 daily sessions of habituation. Habituation was performed by aliplowing the mice to explore the testing chamber for 10 minutes every day. The mice were further habituated to i.p injections for 10 mins every day. One day before the testing day, the mice were moved to dark animal housing, where they were kept until 72 hours after the testing. On the testing day, the mice in the test group were placed in an ANY-maze controlled Fear Conditioning System (Ugo Basile). They were given 20 random shocks (0.5 mA, 2 seconds long) distributed in 10 minutes of testing. The i.p. injection of 4-OHT (10 mg/kg) was performed 2 hours after testing. The control group remained in the home cage but was also habituated to i.p. injections and was injected with 4OHT on the testing day.

### Perfusion and Immunohistochemistry

The animals were anaesthetized with FMM and euthanized by transcardial perfusion with 50 ml of PBS (with 50 mg/ml heparin), followed by perfusion with freshly prepared 4% Paraformaldehyde (PFA). The brains were extracted from the skull and stored in PFA for two days at +4 °C. Brains were sliced into 80um thick slices in PBS on a Leica vibratome.

To enhance the fluorescent signals, an immunohistochemical staining was performed for the eGFP labeling in the odor driven innate fear and shock activated experiments. Antibodies used were an eGFP rabbit (Invitrogen (CAB4211)) primary antibody with 1:1000 dilution and 72-hour incubation. Alexa fluor 488 goat anti-rabbit (Invitrogen (A-11008)) was used as the secondary antibody with 1:1000 dilution with 24-hour incubation. Nuclear staining was done using 1:1000 dilution of DAPI for 30 minutes. Brain slices were mounted on glass slides with coverslips using Fluoromount G Southern Biotech).

### Imaging and cell count

Imaging of the brain slices following the optogenetic stimulation of LA was done using the ZEN software (ZEISS) and a ZEISS Apotome microscope preparing tile images of the whole brain slice. All imaging for the odor driven innate fear and shock-activated tracing experiments was performed using the ZEN software and a ZEISS confocal microscope. The images were taken as z-stack images (z-step size = 2 um) in the respective areas using a x20 objective at the resolution of 1024 pixels X 1024 pixels. Green fluorescence was excited at 488 nm and detected with a bandpass filter of 509-605 nm, while the blue fluorescence was stimulated at 405 nm and detected through a 426-488 nm bandpass filter. In order to avoid photoconversion of mCherry fluorescence, the imaging of the blue channel was always performed after imaging of the red and green channels^4,20^. Cell counting for the optogenetic activation experiment was performed blindly and manually in ImageJ. Cell counting for odor driven innate fear and shock-activated circuit experiments were performed blindly and semi-manually using the cell count function in Imaris (Bitplane).

### Statistics

Statistical analyses were performed in PRISM 6 (GraphPad). Data from male and female subjects were pooled in all experiments apart from the TMT experiment, which only included male subjects. Data were tested for normality with D’Agostino-Pearson normality test, and a parametric test was used for the data that presented a normal distribution. If not, a non-parametric test was performed. Statistical method and corresponding p values are reported in the figure descriptions.

---

**Supp. Fig. S1:**
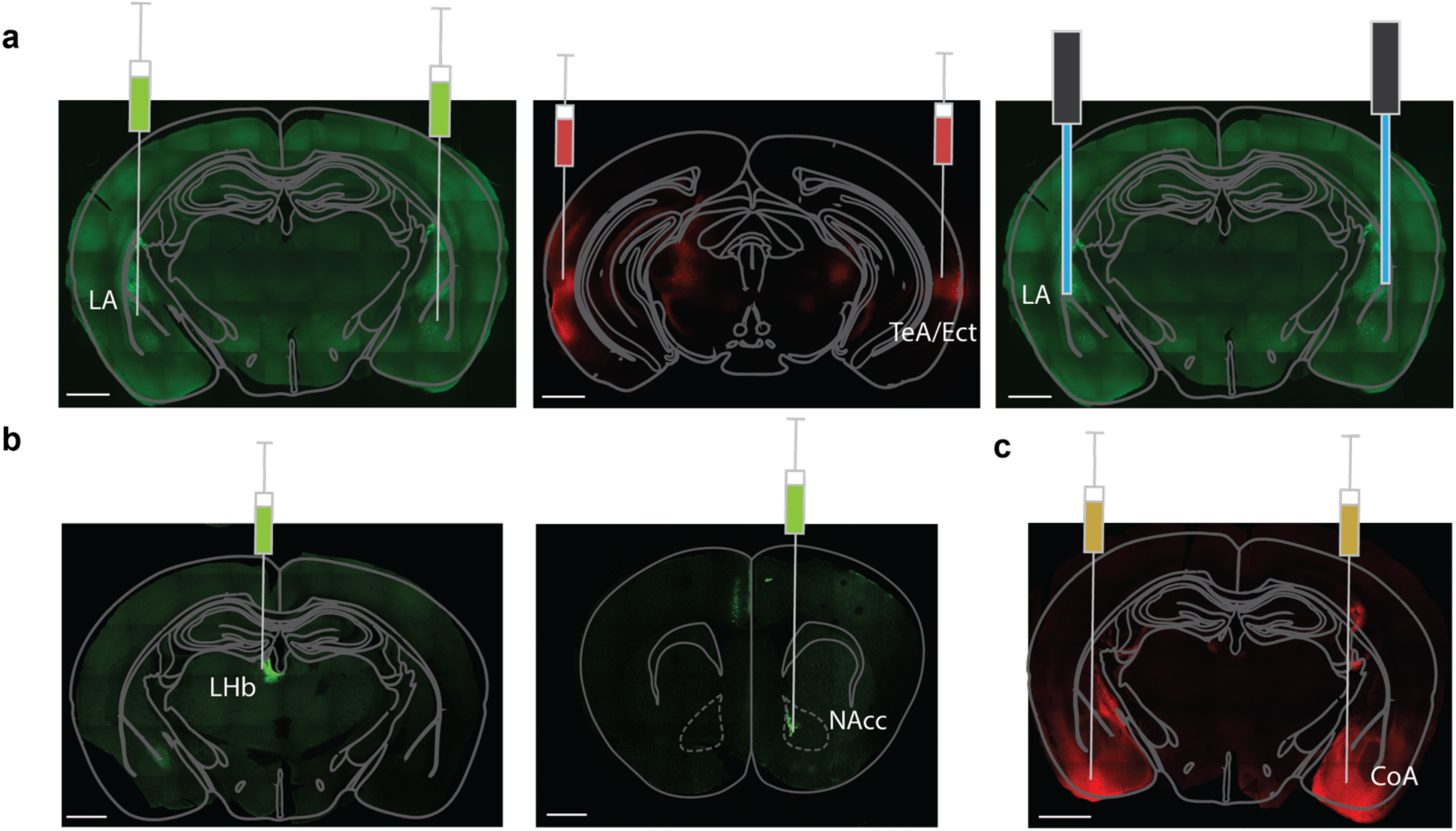
Virus injection sites in mice used in this study. a) Injection site for virus in LA, fiber placement above LA and injection site for AAV-oChIEF-tdTomato in the TeA/Ect. b) Injection site for virus in LHb. Injection site for FluoSpheres and virus in NAcc. c) Injection site for virus in CoA.

**Supp. Fig. S2:**
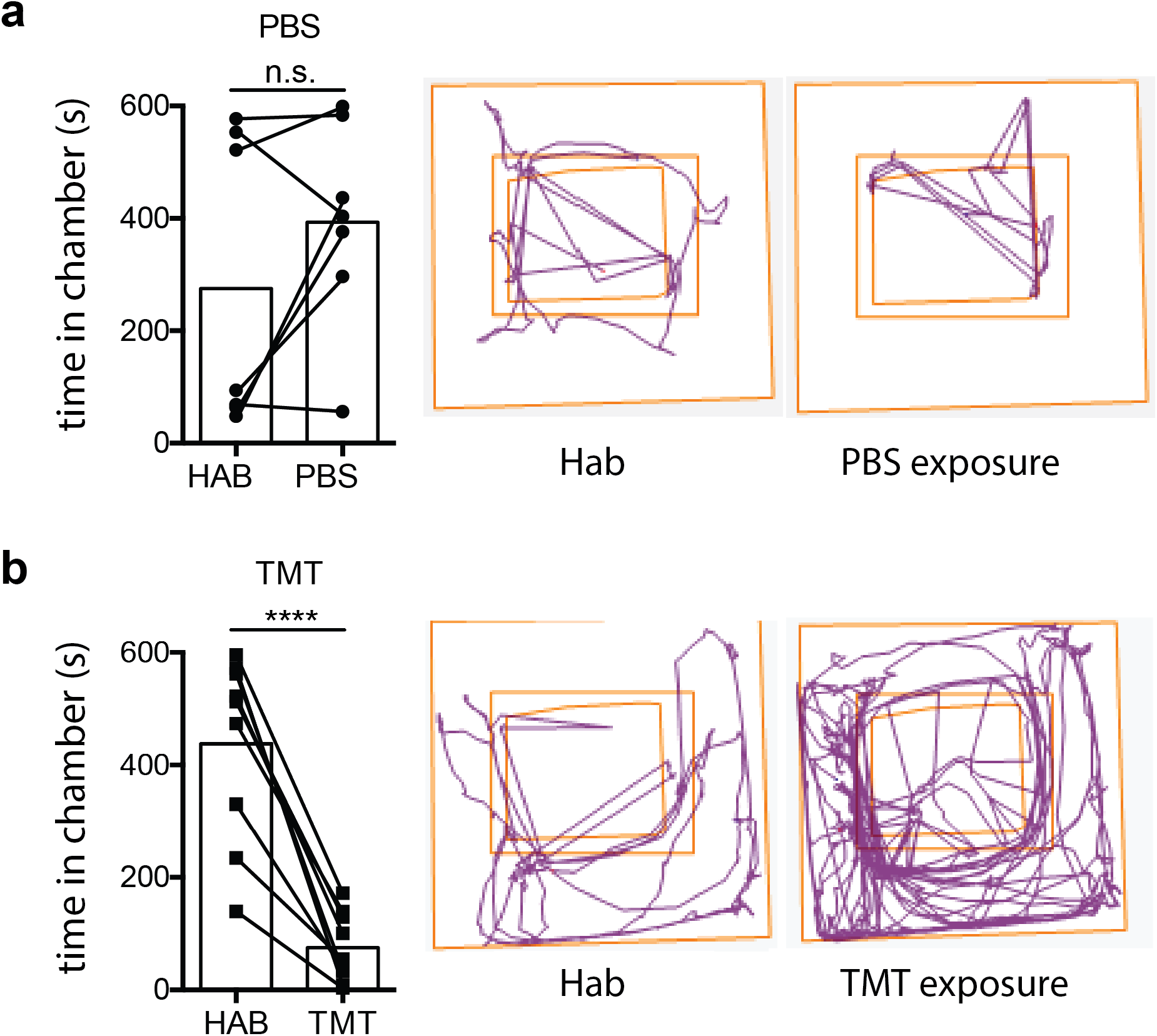
TMT exposure decreases the time spent in the dark chamber and increases the open field exploration. a) Behavioral tracking using ANY-maze. The control group exposed to PBS showed no significant difference in time spent in the dark chamber during the last habituation phase vs. testing. (n = 7; p-value = 0.2188 (Wilcoxon test)). b) Behavioral tracking using ANY-maze. The TMT exposed group showed a significant decrease of time spent in the dark chamber when exposed to TMT in comparison to the last habituation. (n = 9; p-value < 0.0001 (paired t-test)). Not shown: The percentage of time spent inside the box during habituation did not significantly differ between the PBS and TMT exposed groups (PBS group n = 7, TMT group n = 9; p=0.1413 (Mann-Whitney test)).

**Supp. Fig. S3:**
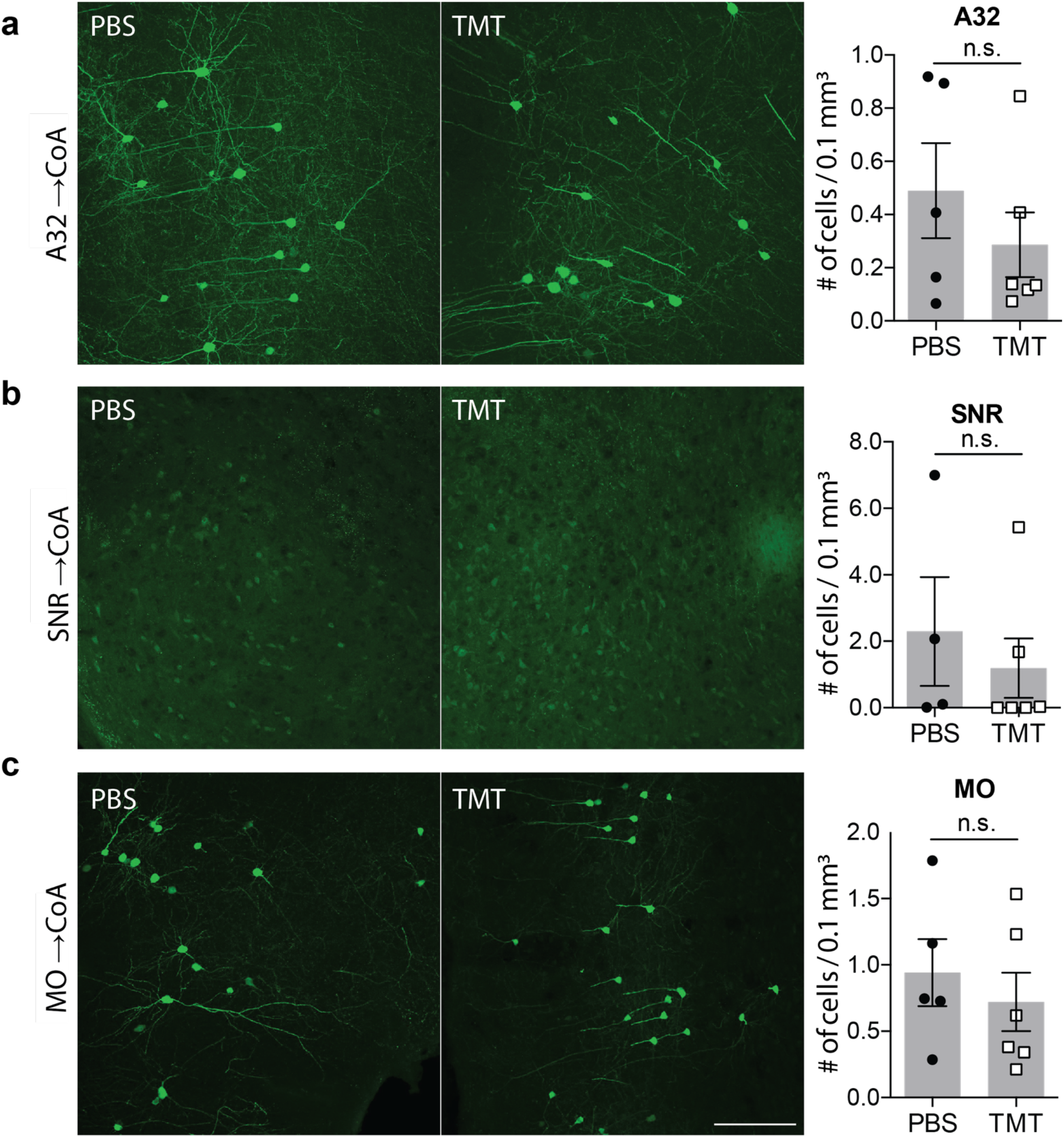
Representative images of the CoA-projecting regions, which deemed insignificant by TRACE in mice exposed to TMT vs PBS. No significant difference of labeling in the CoA-projecting neurons within: a) anterior cingulate cortex area 32 (PBS n = 5, TMT n = 6; p-value = 0.4242 (Mann-Whitney test)), b) substantia nigra reticulata (PBS n = 4, TMT n = 6; p-value = 0.2429 (Mann-Whitney test)), c) medial orbital cortex (PBS n = 5, TMT n = 6; p-value = 0.5281 (Mann-Whitney test))

